# The bat Influenza A virus subtype H18N11 induces nanoscale MHCII clustering upon host cell attachment

**DOI:** 10.1101/2024.07.12.603249

**Authors:** Maria Kaukab Osman, Jonathan Robert, Lukas Broich, Dennis Frank, Robert Grosse, Martin Schwemmle, Antoni Wrobel, Kevin Ciminski, Christian Sieben, Peter Reuther

## Abstract

Prior to the discovery of the bat influenza A virus (IAV) subtypes H17N10 and H18N11, it was believed that all IAVs bind via the viral hemagglutinin (HA) to sialic acid residues to mediate attachment and subsequent viral entry. Both HA subtypes, H17 and H18, however, do not bind to sialic acids but instead engage a proteinaceous receptor: major histocompatibility complex class II (MHCII). The mechanistic details of this hitherto unknown protein-mediated entry are not understood. Since conventional IAVs require attachment to clusters of sialylated glycans to overcome the low affinity of the HA-sialic acid interaction, we hypothesized that bat HA would likewise interact with multiple MHCII molecules. Here, we used photoactivated localization microscopy (PALM) on fixed and live cells expressing MHCII fused to appropriate fluorescent reporters. We show that bat IAV particles attach to pre-existing MHCII clusters present on susceptible cells and that the local HA-MHCII interaction results in an increased cluster size. To measure the impact of viral attachment on the dynamics of MHCII, we utilized an experimental setup designated “inverse infection” where intact viral particles were immobilized on coverslips before live MHCII-expressing cells were seeded on top. Recording the trajectories of single MHCII molecules, this approach revealed that the mobility of MHCII was indeed slowed down in viral proximity leading to a local enrichment of MHCII molecules beneath the viral particle. Taken together, these data suggest that attachment of viral particles leads to clustering of MHCII, a process similar to the MHCII dynamics observed during the formation of an immunological synapse.

## Introduction

Influenza A viruses (IAV) have a major impact on global health causing annual epidemics and sporadic pandemics. Until 2012, aquatic birds were believed to represent the sole reservoir of IAV maintaining all previously known hemagglutinin (HA) (H1-16) and neuraminidase (NA) (N1-9) subtypes^1–3^. This notion changed with the discovery of two novel IAV subtypes, designated H17N10 and H18N11, in bats from Central and South America^4–6^. While these bat IAVs structurally resemble conventional IAVs of avian origin, their HA and NA surface glycoproteins are functionally distinct. In contrast to conventional HAs, the bat IAV HAs (H17 and H18) do not bind sialic acids for cell entry and the correspondent NAs (N10 and N11) lack sialidase activity^5,7–9^. Previous studies using recombinant vesicular stomatitis virus (VSV) expressing either H18 or N11, have shown that H18 but not N11 is sufficient to mediate cell entry and allow viral spread^10^. Consistent with these observations, we demonstrated that a mutant H18N11 lacking the N11 ectodomain (designated rP11) not only replicates efficiently in cell culture but also in its natural host the Jamaican fruit bat (*Artibeus jamaicensis*)^11^. Even though N11 is still required for efficient transmission among bats, H18 thus seems to be the key determinant of effective infection at the cellular level. However, the receptor(s) involved in H18-mediated entry remained unknown until recently.

We and others showed that bat IAVs rely on a proteinaceous receptor for cell entry: the major histocompatibility complex class II (MHCII)^12,13^. MHCII is a heterodimeric transmembrane protein complex consisting of an α and β chain, each comprising two extracellular domains: α1, α2 and β1, β2^14^. MHCII is mainly expressed on professional antigen-presenting cells (APC) such as macrophages, dendritic cells and B cells^15^. Here, MHCII has an essential role in adaptive immunity by presenting peptides from endocytic compartments to CD4^+^ T cells. Interestingly, MHCII from various vertebrate species including the human leukocyte antigen DR (HLA-DR), support H17 and H18-mediated infection and the highly conserved amino acid residues within the α2 domain of MHCII are required for cell entry^12,16^. However, the initial steps of bat IAV infection, including receptor engagement, endocytosis and endosomal release, remain elusive. In-depth analysis of receptor engagement by classical biochemical approaches has been unsuccessful likely due to a low affinity between bat HA and MHCII, a feature reminiscent of the HAs of conventional IAVs, which bind sialic acids very weakly^17–19^. So far, an MHCII-H18 interaction could only be confirmed by chemical crosslinking on the cell surface^12^.

Conventional IAV particles interact with sialylated cell surface glycoproteins^20^, which are organized in submicrometer nanoclusters^21,22^. These clusters represent multivalent virus binding platforms that provide the avidity necessary for attachment of multiple low-affinity HA’s and subsequent endocytosis of the viral particle. Based on a previous observation that MHCII is enriched in membrane clusters of APCs, we hypothesized that these MHCII clusters also serve as multivalent attachment sites^23–25^. As our recent *in silico* model suggests that one H18 homotrimer can bind three MHCIIs, we speculate that upon attachment of bat IAV particles, additional MHCII complexes are recruited^16^.

Here, we use photoactivated localization microscopy (PALM) to visualize the nanoscale organization of MHCII and the interaction dynamics of bat IAV and MHCII in live cells at the single-molecule level^26,27^. We show that individual bat IAV particles interact with clusters of MHCII, resulting in decreased mobility of the viral particle at the cell surface. Using an “inverse infection” approach, we show that additional MHCII molecules are trapped at the virus-cell interface, resulting in increased MHCII cluster size, suggesting that viral particles induce nanoscale MHCII clustering upon host cell attachment.

## Results

### Cells expressing MHCII fused to the photoconvertible fluorescent protein mEos3.2 are highly susceptible to infection with the bat IAV H18N11

To visualize the dynamics of MHCII upon attachment of H18N11, we transduced non-permissive MDCK-II cells with a DNA cassette from which the wildtype (wt) alpha and beta chain of the human MHCII HLA-DR15 are expressed (Fig 1A). As we have shown recently that covalent fusion of a model peptide to MHCII enhances the susceptibility to bat IAV infection, the antigenic HA_307-319_ peptide was linked to the extracellular N-terminus of the beta chain^16^. To allow visualization of MHCII at the single-molecule level in live cells by PALM, we fused the photoconvertible fluorescent protein mEos3.2 to the intracellular C-terminus of the beta chain (MHCII_mEos_)^28^. In the native state, mEos3.2 emits green fluorescence, but can be irreversibly converted by UV-illumination at 405 nm. Photo-converted mEos3.2 emits red fluorescence when excited with a 561 nm laser before it eventually photobleaches (Fig. 1B, Fig. S1A). In PALM, this photoswitching mechanism is used to localize individual mEos3.2 molecules over time allowing reconstructing of the organization and dynamics of single MHCII molecules on the cell surface. We also generated a mutant MHCII (MHCII_mEosmut_) with 11 amino acid substitutions in the α2 subunit derived from the non-classical MHCII, HLA-DM (Fig. 1A), which we recently demonstrated to not support bat IAV infection^16^. Both MHCII_mEos_ and MHCII_mEosmut_ were efficiently expressed at the cell surface of transduced MDCK cells (>99%) (Fig. 1C). Furthermore, both wt and mutant MHCII, here fused to the red-fluorescent TagRFP (MHCII_TagRFP_ and MHCII_TagRFPmut_), activated T cells in a haplotype-specific manner confirming the overall structural integrity of the proteins (Fig. 1D, Fig. S1 B-D).

**Figure 1:**
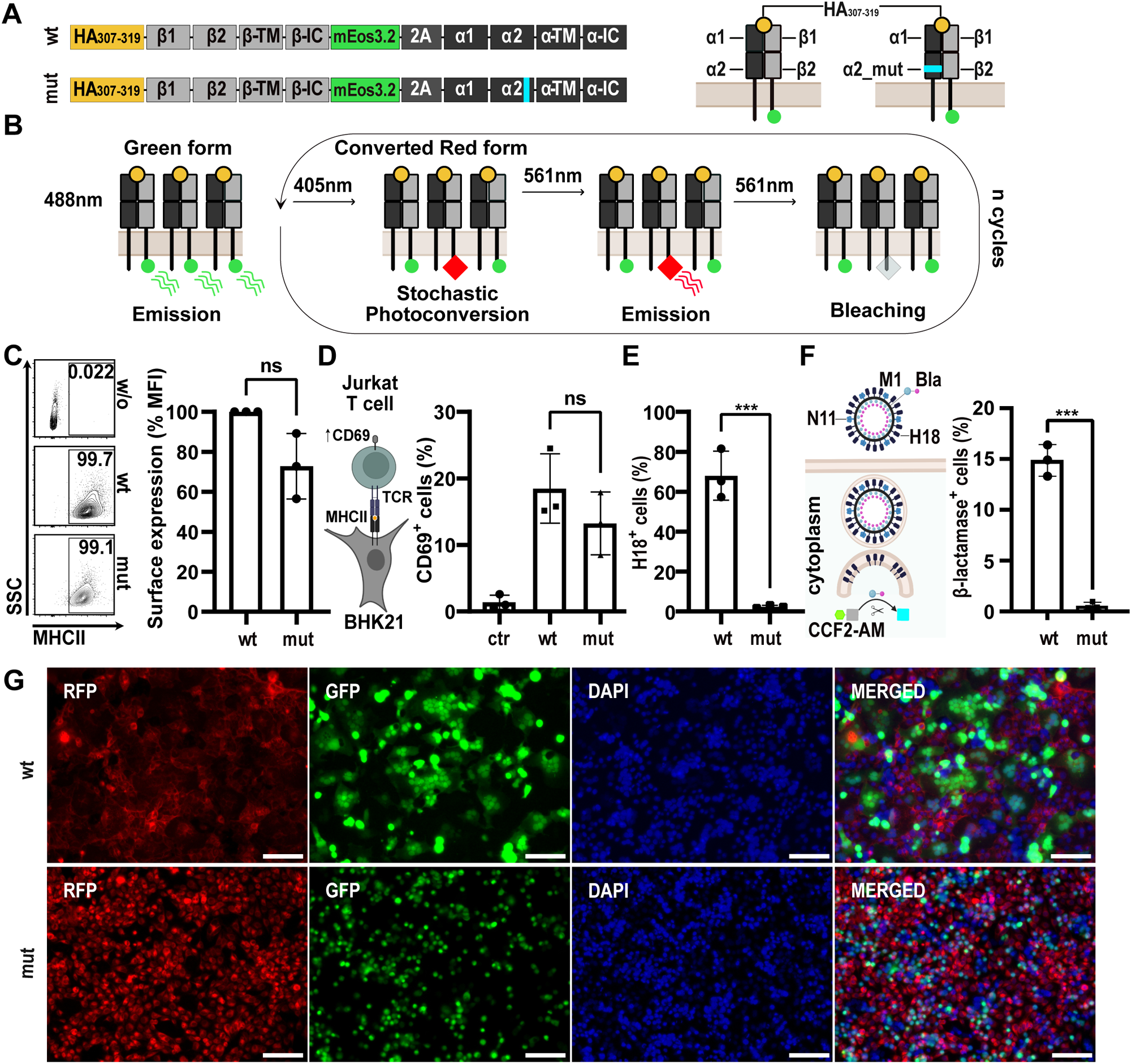
Fluorescently-labeled MHCII mediates entry of bat IAV. **(A)** Illustration of the fluorescently-labeled human MHCII (HLA-DR) constructs used for this study: MHCII_mEos_ and MHCII_mEosmut_. The extracellular N-terminus of the beta chain is fused to the influenza HA_307-319_ model peptide and the intracellular C-terminus is linked to the photoconvertible mEos3.2 protein. The α2 subunit of MHCII_mEosmut_ comprises a substitution of 11 amino acids (cyan), which are derived from the non-classical MHCII HLA-DM. Co-translational separation of the two MHCII chains is mediated by a PTV-1 2A sequence **(B)** Schematic representation of the photoconversion of mEos3.2. Upon 405 nm light exposure, single mEos3.2 fluorophores convert stochastically. Converted mEos3.2 emits red light upon 561 nm excitation and then eventually photobleaches. For PALM, red fluorescence is recorded over 10,000 frames. **(C)** Flow cytometric analysis of the surface expression of MHCII_mEos_ (wt) and MHCII_mEosmut_ (mut) in transduced MDCK-II cells. Small contour plots illustrate MHCII surface expression levels. Numbers within the plots indicate the percentages of MHCII-positive cells. The bar graph depicts the median fluorescent intensity of the MHCII signal normalized to wildtype **(D)** Cartoon depicting the principle of the T cell activation assay. BHK21 cells transiently expressing MHCII_TagRFP_ (Haplotype: HLA-DR1), are co-cultured with CH7C17Jurkat T cells. These T cells are transgenic for the HA1.7 T cell receptor (TCR), which recognizes the HA_307-319_ when presented by HLA-DR1. Activated T cells express CD69 on their surface. The bar graph shows the number of CD69 positive T cells upon activation by MHCII_TagRFP_ (wt) or MHCII_TagRFPmut_ (mut). Co-culture of T cells with MHCII_TagRFP_ of the haplotype HLA-DR15 served as negative control (ctr.). **(E)** Infection rate of MDCK-II cells stably expressing MHCII_mEos_ (wt) or MHCII_mEosmut_ (mut) at 24 hours post-infection with H18N11 at an MOI of 5. Percentage of infected cells was determined by flow cytometric analysis of the H18 expression. **(F)** Quantification of viral entry using H18N11 VLPs harboring the viral matrixprotein (M1) fused to a β-lactamase (M1-Bla). Upon endosomal escape, β-lactamase activity in the cytoplasm was determined with the FRET-based reporter CCF2-AM. CCF2-AM consists of blue-fluorescent hydroxycoumarin and green-fluorescent fluorescein linked via a β-lactam ring. CCF2-AM emits green fluorescence via FRET when excited at 409 nm. β-lactamase separates the two fluorophores resulting in blue fluorescence at 409 nm, here measured by flow cytometry. The bar graph depicts the percentage of β-lactamase positive MDCK-II cells expressing MHCII_TagRFP_ (wt) or MHCII_TagRFPmut_ (mut) upon exposure to the VLPs. **(G)** pH-induced polykaryon formation of HEK293T cells expressing H18 and GFP with MDCK-II cells stably expressing MHCII_TagRFP_ (wt) or MHCII_TagRFPmut_ (mut). Images are representatives of 3 independent experiments. Scale bar 100 µm. Error bars represent SD. Paired Student’s t-test was applied for (C) and unpaired two-tailed Student’s t test was used in (D-F). *** P < 0.001, ns: not significant. (D) and (F) Schemes were created with BioRender.com

Next, we compared the ability of each of the two MHCII constructs to support bat IAV infection. While expression of MHCII_mEos_ rendered cells susceptible to infection with the H18N11 variant rP11, MHCII_mEosmut_ did not (Fig. 1E, Fig S1C-D). Using H18N11 virus-like particles (VLPs) harboring an M1-β-lactamase (M1-Bla) fusion protein, we further probed at which stage the infection is blocked^12^. We detected no cytoplasmic Bla activity in the MHCII_mut_-expressing cells proving that no viral release from the endosome occurred (Fig 1F). In addition, H18-mediated cell fusion was only observed in presence of the wt but not the mutant form, demonstrating that H18 is unable use mutant MHCII for membrane fusion (Fig. 1G, Fig. S1E).

To test whether the incompatibility of MHCII_mEosmut_ with H18N11 infection is due to an insufficient interaction on the cell surface, we produced soluble ectodomains of wt MHCII (sMHCII) and mutant MHCII (sMHCII_mut_) and tested their ability to neutralize H18N11 (Fig. 2A-C). sMHCII and sMHCII_mut_ were purified from the supernatant of transfected Expi293F cells and incubated with H18N11 prior to infection of MDCK-II cells stably expressing MHCII_mEos_. Under these conditions, sMHCII efficiently neutralized H18N11 particles in a dose-dependent manner, as determined by counting the number of infectious viruses (focus forming units) left after treatment (Fig. 2D). No virus neutralization was observed for sMHCII_mut_.

**Figure 2:**
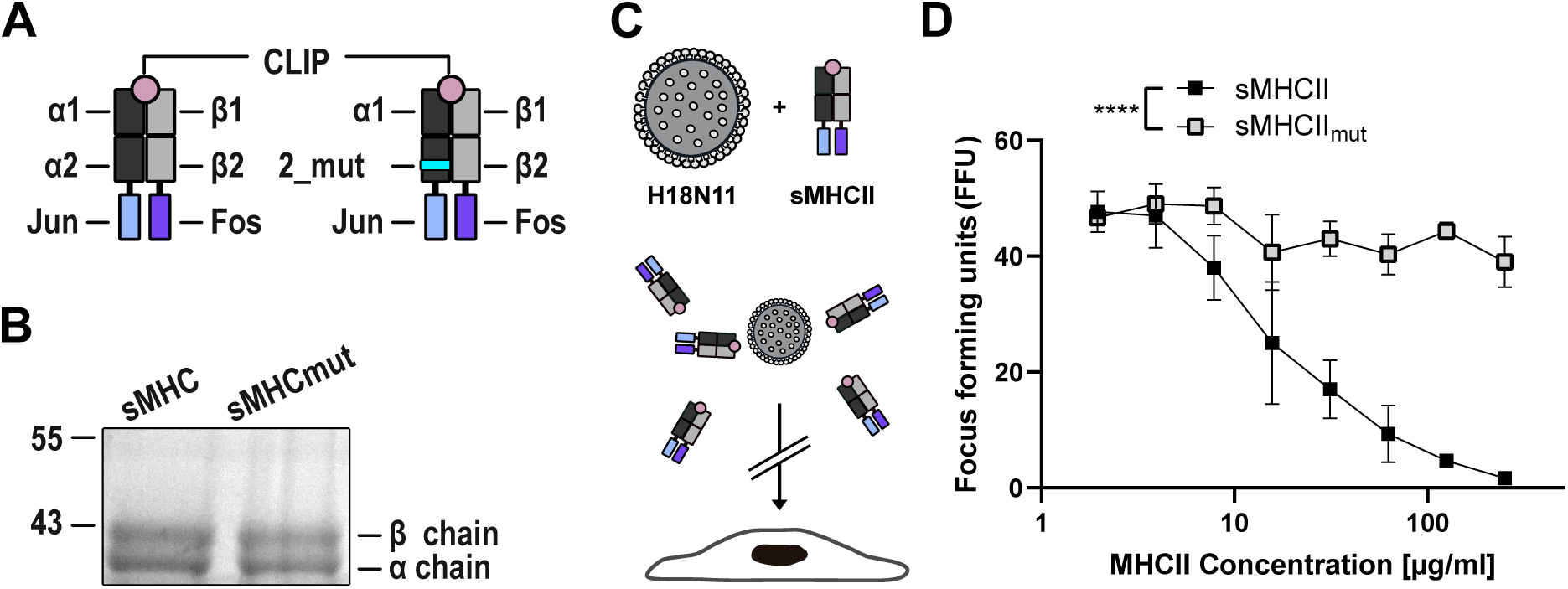
Soluble, wildtype MHCII efficiently neutralizes H18N11 viral particle. **(A)** Illustration of soluble MHCII (sMHCII) and mutant (sMHCII_mut_). The C-termini of the α and the β chain were fused to the zipper motifs Jun and Fos, respectively. The N-terminus of the β-chain is fused to the CLIP peptide. The 11 amino acid substitution within the sMHCII_mut_ α-chain is highlighted in cyan. **(B)** Coomassie staining of the purified sMHCII and sMHCII_mut_. **(C)** Schematic illustration of the H18N11:MHCII competition assay. H18N11 viral particles are incubated in presence of soluble MHCII prior to infection. **(D)** Comparison of the ability of the indicated concentrations of sMHC and sMHC_mut_ to neutralize 50 focus-forming units of H18N11. Error bars represent SD. Two-way ANOVA was used for statistical analysis, **** p < 0.0001. (A and C) Schemes were created with BioRender.com

In summary, our fluorescently-labeled wt MHCII supports bat IAV infection and is therefore suitable for further functional studies using high-resolution microscopy. Cell surface-expressed MHCII_mEosmut_ will serve as an adequate negative control, since it does not support viral infection, but is still able to activate T cells.

### H18N11 particles bind to MHCII clusters on the cell surface

Since conventional IAVs overcome their poor affinity to sialic acid by multivalent binding to clusters of sialylated glycans on the cell surface^21,29^, we hypothesized that H18N11 may preferentially bind to preformed clusters of MHCII. To visualize the molecular organization of MHCII on the cell surface, MDCK-II cells expressing either MHCII_mEos_ or MHCII_mEosmut_ were fixed and imaged with PALM (Fig. 3A). We observed clusters on the cell surface with a comparable median radius of gyration (Rg) of 33 nm for wt MHCII and 40 nm for mutant MHCII (Fig. 3 B-C, Fig S2A). This range of cluster sizes is in agreement with the previously observed organization of MHCII expressed endogenously on professional APCs^30^. Next, we set out to determine the size of MHCII clusters interacting with H18N11 viral particles. Cells were exposed to the H18N11 variant rP11^11^ on ice to allow attachment to the cell surface but without subsequent internalization of viral particles. PALM revealed MHCII_mEos_ co-localizing with single viral particles after fixation and anti-H18 immunolabeling. We found that clusters underneath viral particles are larger with a median Rg of 48 nm, compared to the median cluster size of Rg = 29 nm at virus-free surfaces (Fig 3D-E). In sharp contrast, the median cluster size of MHCII_mEosmut_ was found to be 37 nm under viral particles (Figure 3F-G). Taken together, viral particles bind to MHCII_mEos_ with increased cluster size on ice, suggesting a direct interaction of H18 viruses with pre-existing MHCII clusters.

**Figure 3:**
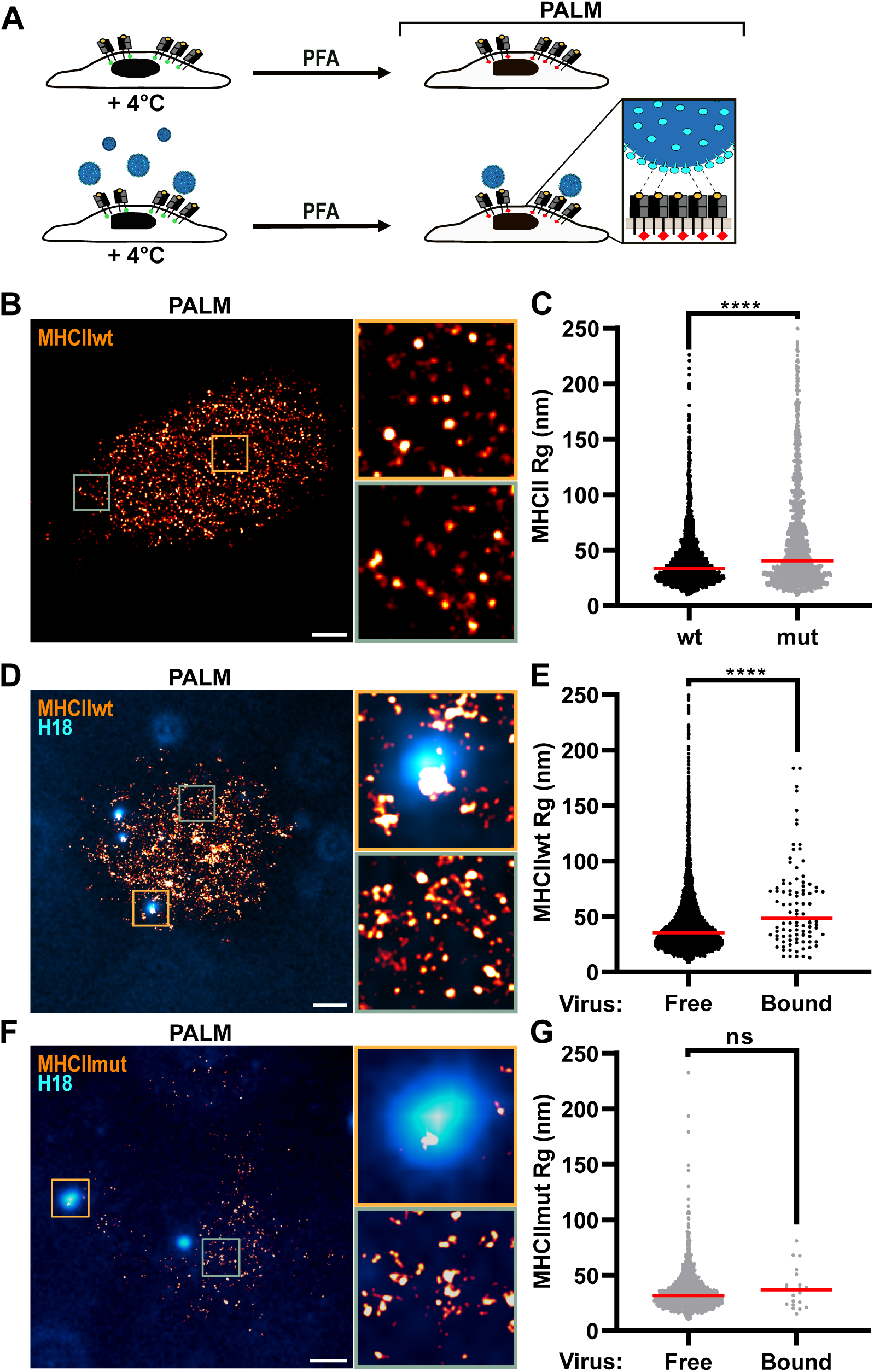
Viral particles are associated with large clusters of wt MHCII but not mutant MHCII. **(A)** Schematic representation of the analysis of the MHCII-distribution at the surface of fixed MDCK-II cells stably expressing mEos-fused MHCII by Photoactivated Localization Microscopy (PALM) in the presence or absence of H18N11 particles. **(B)** MHCII-distribution on the surface of noninfected MDCK-II cells stably expressing MHCII_mEos_ (wt) by PALM **(C)** Comparison of MHCII cluster sizes at the surface of noninfected MDCK-II cells stably expressing MHCII_mEos_ (wt) and MHCII_mEosmut_ (mut). **(D)** MHCII-distribution at the surface of MDCK-II cells stably expressing MHCII_mEos_ (wt) in the presence of H18N11 viral particles determined by PALM. Viral particles were visualized by staining for H18 (cyan). **(E)** Comparison of MHCII cluster sizes at virus-free and virus-bound surfaces of MDCK-II cells stably expressing MHCII_mEos_ (wt) in the presence of H18N11 viral particles. Data was pooled from at least two independent experiments. **(F)** MHCII-distribution at the surface of MDCK-II cells stably expressing MHCII_mEosmut_ (mut) in the presence of H18N11 viral particles. **(G)** Comparison of MHCII cluster sizes at virus-free and virus-bound surfaces of MDCK-II cells stably expressing MHCII_mEosmut_ (mut) in the presence of H18N11 viral particles. Data was pooled from at least two independent experiments. Unpaired, two-tailed Student’s t-test was used for comparison. **** P < 0.0001, ns: not significant. (A) Schemes were created with BioRender.com

### Interaction between H18 viruses and MHCII receptors decreases the mobility of both, the viral particles and MHCII

For conventional IAV, we have recently shown that viral particles become transiently confined to the host cell surface when they encounter a region of high receptor density^21^. Similarly, we hypothesized for H18N11 particles that binding of MHCII may decrease both, the viral mobility and the local receptor diffusion leading to larger clusters as observed in fixed cells (Fig. 3D and E). To visualize the dynamic movement of H18N11 on the cell surface, particles were fluorescently labeled and viral trajectories were recorded by spinning-disk confocal microscopy (Fig. 4A). For this purpose, MHCII-expressing cells were incubated with DiO-labelled H18 particles at 4°C to allow virus-cell binding while preventing uptake. Subsequently, cells were transferred to 37°C and trajectories were recorded over 20 min. This analysis revealed short viral trajectories on cells expressing MHCII_mEos_, whereas trajectories were significantly longer on MHCII_mEosmut_ cells (Fig. 4B). This suggests an MHCII-dependent confinement of the H18N11 particle on the cell surface.

**Figure 4:**
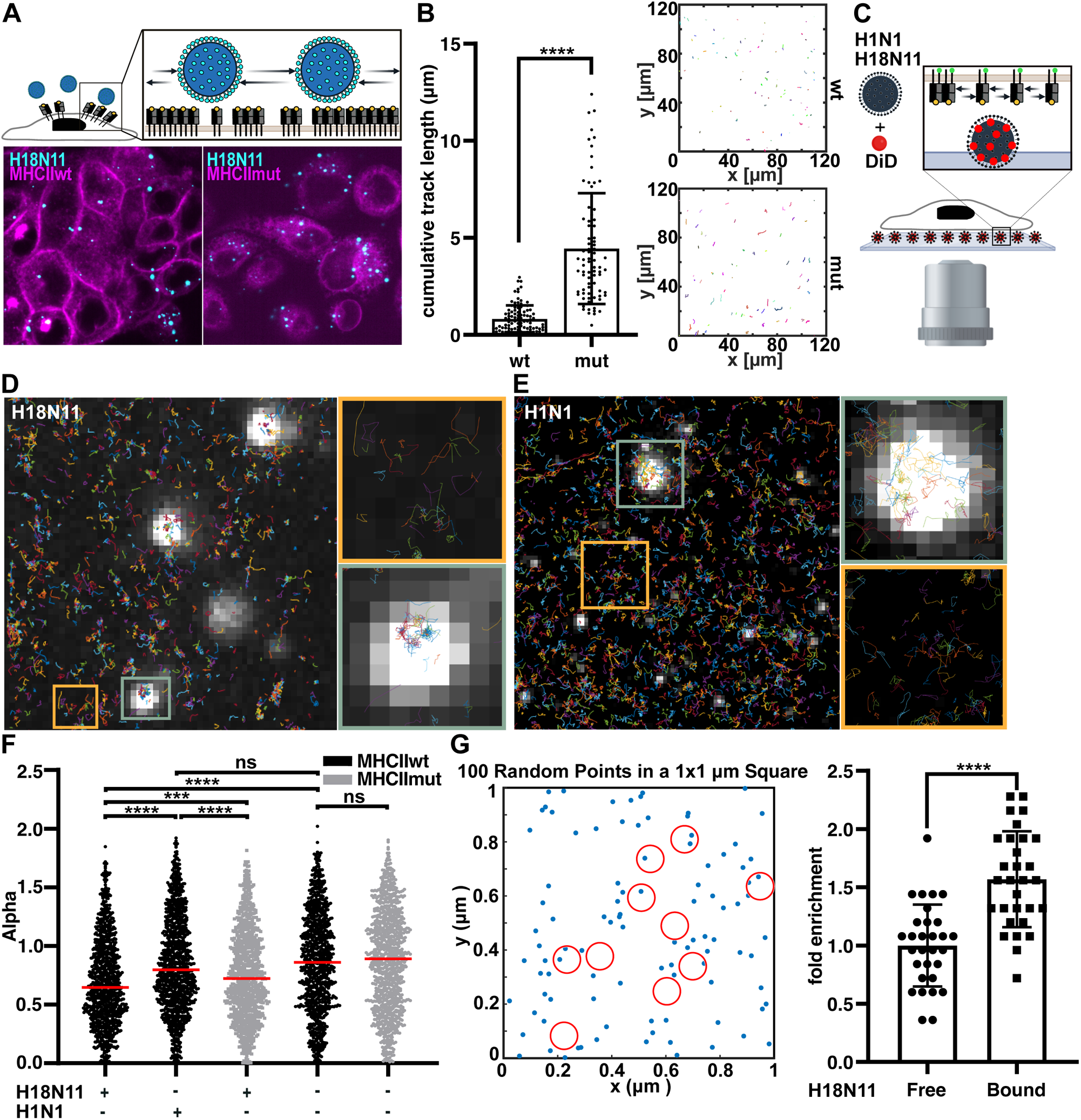
MHCII is enriched underneath H18N11 viral particle upon attachment. **(A)** Schematic representation of the tracking of viral particles on the cell surface (upper panel). Here, fluorescently-labeled viral particles were added to MDCK-II cells stably expressing MHCII_TagRFP_ or MHCII_TagRFPmut_ (lower panel). **(B)** Representative trajectories of fluorescently labeled H18N11 viral particles on MDCK-II cells stably expressing MHCII_TagRFP_ or MHCII_TagRFPmut_ (right) and comparison of the cumulative track length (left). **(C)** Schematic illustration of the inverse infection assay. Fluorescently labeled H18N11 viral particles were immobilized on epoxy-modified glass slides. MDCK-II cells stably expressing MHCII_mEos_ or MHCII_mEosmut_ were seeded on top. **(D and E)** Tracking of MHCII_mEos_ molecules by live-PALM on MDCK-II cells stably expressing MHCII_mEos_ seeded on top of immobilized H18N11 (D) or H1N1 (E) viral particles, which are depicted in white. A representative area including a viral particle and a virus-free area is shown at a higher magnification. **(F)** Comparison of the power-law exponent α of the respective MHCII molecules on the surface of transduced MDCK-II cells exposed to the indicated viruses at virus-bound areas. Mock-infected MHCII_mEos_ or MHCII_mEosmut_ cells surfed as control. **(G)** To calculate if a slowdown of receptor proteins at the virus binding site leads to local protein enrichment, we simulated the 2D random diffusion of MHCII using our measured diffusion coefficient D_MHCII_ = 0.032 µm^2^/s. We then added circular regions (red, r = 50 nm) to the simulation, where the proteins were slowed down according to our PALM measurements to D_MHCII+H18_ = 0.015 µm^2^/s. We simulated 4000 time steps with 30 ms interval. At the end, we examined the number of receptors associated with the circular regions (i.e., viruses). (F and G) Unpaired two-tailed Student’s t-test was used for pairwise comparison. **** P < 0.00001***, P < 0.0001, ns: not significant. (A and C) Schemes were created with BioRender.com

We further speculated that the H18:MHCII interaction does not only reduce the mobility of the viral particle, but that viral attachment also spatially confines MHCII. Tracking individual MHCII molecules relative to the viral particle by live-cell PALM at physiological temperature, however, is challenging, as viral particles are rapidly internalized. To circumvent this problem, we established a novel experimental setup designated “inverse infection” (Fig. 4C, Broich et al., 2024). Here, fluorescently-labeled viral particles were covalently immobilized on reactive epoxy-modified glass slides and MDCK-II cells expressing MHCII_mEos_ or MHCII_mEosmut_ were seeded on top. Live-cell single-particle tracking (spt)PALM revealed that MHCII_mEos_ molecules diffuse freely in virus-free surface areas, while their tracks were more condensed above the viral particles (Fig. 4D, Fig. S2B). Notably, we did not observe this degree of immobilization of MHCII with conventional IAV A/PR/8/34 (H1N1) suggesting a virus-specific interaction (Fig. 4E). To better quantify the local confinement, we calculated the mean squared displacement (MSD) of all measured MHCII trajectories. An MSD vs. lag time plot can be fitted to a power-law distribution (see methods) where the power-law exponent (α) can be used as a measure for local confinement^31^. While α of 1 indicates free diffusion, smaller α values suggest a restricted lateral mobility. As shown in Figure 4F, the contact with an H18N11 particle resulted in local confinement of MHCII_mEos_ (median α *=* 0.64), whereas the mobility of MHCII_mEos_ in proximity of an H1N1 particle was almost not restricted (median α *=* 0.8). The local confinement of MHCII_mEos_ in contact with H18N11 particle was significantly higher than that of MHCII_mEosmut_. Yet, for MHCII_mEosmut_, some degree of immobilization above the viral particle was observed compared to uninfected cells suggesting a residual affinity of the mutant MHCII to H18 (Fig. 4F). Virus-induced receptor confinement as observed here should lead to local enrichment of MHCII proteins in proximity of the virus. To show this, we simulated a patch of the plasma membrane with diffusing MHCII receptors. We then added circular regions (i.e. bound viruses) where the receptor diffusion was reduced due to virus-receptor interaction (Fig. 4G). The corresponding diffusion coefficients (D_MHCII_ and D_MHCII+H18_) were taken from the power-law exponent analysis described above (see methods). We then simulated 4000 time steps (interval 30 ms) and counted the number of receptors in viral proximity. For the H18-mediated diffusion slowdown measured for MHCII, we indeed found a significant local enrichment as compared to freely diffusing receptors (Fig. 4G).

In summary, we show that the interaction between H18 and MHCII results in decreased viral mobility on the host cell surface and local confinement of individual receptors in viral proximity. The H18:MHCII interaction slows down MHCII diffusion which leads to a characteristic local enrichment leading to larger MHCII clusters beneath the viral particles (as shown in Fig. 3D and E).

## Discussion

The discovery of influenza A viruses in New World bats changed the paradigm that entry of IAV is strictly glycan-dependent. While MHCII was identified as the primary receptor, detailed insights into the entry of bat IAV have been missing. Here, we reveal that viral attachment to the host cell surface via MHCII is a highly dynamic process. We show that H18N11 associates with large MHCII clusters resulting in a local confinement of the viral particle. This is consistent with observations from conventional IAV, where multivalent receptor-ligand interactions are necessary to decrease the overall dissociation rate^21,22^. In contrast to conventional IAVs, bat IAVs, do not attach to nanodomains composed of diverse sialylated glycans, but rely on a single proteinaceous receptor. Consistently, the mutant MHCII used in this study, which carries mutations in the proposed H18 binding site, does not support viral entry. Using the “inverse infection” approach (Broich et al., 2024), we can show that the interaction of the bat IAV particle with MHCII also results in a reduced mobility of MHCII. Interestingly, it was demonstrated that cross-linking of MHCII by natural ligands or MHCII-specific antibodies induces downstream signaling, which promotes clathrin-dependent endocytosis^32–35^. It is therefore tempting to speculate that attachment of H18N11 similarly triggers intracellular signaling resulting in the uptake of MHCII:H18N11 complexes. Likewise, outside-in signaling by receptor tyrosine kinases such as EGFR is also required for the efficient uptake of conventional IAVs^36^. Thus, despite of the fundamentally different nature of the cellular receptor, the early events of the bat IAV entry seem to resemble that of conventional IAVs.

Subsequent to endocytosis, the IAV envelope has to fuse with the endosomal membrane^37^. Performing cell fusion assays, we could prove that viral-host membrane fusion is also dependent on the interaction between H18 and MHCII^12,16^. Accordingly, the mutant MHCII, which cannot be enriched under the viral particle on the cell surface does not promote membrane fusion. Based on this finding, we hypothesize that for endosomal escape, viral particles also require a multivalent interaction with MHCII inside the endosome.

The selective pressure that drove viral evolution towards the use of a proteinaceous receptor instead of sialic acid is still unclear, but illustrates that IAVs have the potential to switch their receptor specificity with unforeseen consequences such as a complete change in cellular tropism^38,39^. Of note, a novel IAV subtype (H19), which was identified recently in an avian reservoir, was shown to also depend on MHCII for cell entry^40^. This demonstrates that MHCII-binding is not a unique feature of bat-borne IAV and might be evolutionarily older than anticipated.

In this study, we shed light on the initial steps of the bat IAV H18N11 replication cycle. We show that viral attachment and subsequent uptake depends on a dynamic interaction between H18 on the viral particle and MHCII on the host cell surface. This interaction results in a confinement of both the virus and MHCII, which seems to be critical for the induction of an intracellular signaling cascade that promotes endocytosis.

## Acknowledgments

We thank Wolfgang Schamel for providing the CH7C17 Jurkat T cells. We are grateful to Silke Stertz for the m1-Bla expression plasmid.

## Funding

This work was supported by grants from the European Research Council (ERC) to M.S. (NUMBER 882631—Bat Flu) and in part by the Excellence Initiative of the German Research Foundation (GSC-4, Spemann Graduate School) and the Ministry for Science, Research and Arts of the State of Baden-Wuerttemberg. M.K.O and J.R. are members of the Spemann Graduate School of Biology and Medicine (SGBM). A.G.W. is supported by the Francis Crick Institute which receives its core funding from Cancer Research UK (FC001078), the UK Medical Research Council (FC001078), and the Wellcome Trust (FC001078). P.R. is supported by the Hans A. Krebs Medical Scientist Programme of the Medical Faculty of the University of Freiburg. C.S. acknowledges support by the Helmholtz Association (VH-NG-1526). The funders had no role in study design, data collection and analysis, decision to publish, or preparation of the manuscript. The Lighthouse Core Facility is funded in part by the Medical Faculty, University of Freiburg (Project Numbers 2023/A2-Fol; 2021/A2-Fol; 2021/B3-Fol) and the DFG (Project Number 450392965).

## Materials and methods

### Cell lines

Human embryonic kidney cells HEK293T cells were obtained from the American Type Culture Collection (ATCC;CRL3216). Baby Hamster Kidney Fibroblasts (BHK-21) cells were obtained from the German Cell Culture Collection (DSZM). MDCK-II cells stably expressing MHCII were generated by lentiviral transduction as described^12^. Two days post transduction cells were selected with 2.5 μg/ml puromycin. Puromycin-resistant cells were single-sorted to obtain clonal cell lines. All adherent cells were cultured in Dulbecco’s Modified Eagle’s Medium (DMEM; Gibco, Thermo Fischer Scientific) containing 10% fetal calf serum (FCS), 100 U/ml penicillin, and 100 mg/ml streptomycin at 37 °C with 5% CO_2_. CH7C17 Jurkat T cells were cultured in Roswell Park Memorial Institute 1640 Medium (RPMI 1640, Gibco, Thermo Fischer Scientific) supplemented with 10% FCS and 5% HEPES at 37°C with 5% CO_2_. Expi293F were obtained from Thermo Fisher and cultured in FreeStyle 293 Expression Medium (Thermo Fisher Scientific) under agitation at 37°C with 8% CO_2_.

### Generation of recombinant bat influenza H18N11 viruses

To generate recombinant cell-culture adapted H18N11 (rP11), the pHW2000-based rescue system was used as described previously^11^. Rescued virus was amplified on MDCK-II cells stably expressing MHCII_mEos_.

### Plasmids

To generate the MHCII_mEos_ plasmid for lentiviral transduction, sequences encoding a signal peptide (MKSLSLLLAVALGLATAVSAGPAV), HLA-DRB1*1501 (NM_001243965.1), mEos3.2 and HLA-DRA (NM_019111.4) were amplified and assembled by overlapping fusion PCR. The sequences encoding the influenza model peptide HA307-319 (PKYVKQNTLKLAT) and the PTV-1 2A peptide (ATNFSLLKQAGDVEENPGP) were introduced as overlapping primer overhangs. The assembled construct was cloned into the pLVX-puro vector using BamHI and EcoRI restriction sites. The sequence encoding the 11 amino acid substitution (EIDRYTAIAYW) resulting in MHCII_mut_ was introduced via overlapping fusion PCR. Plasmids encoding MHCII_TagRFP_ and MHCII_TagRFPmut_ for lentiviral transduction were generated analogously. For transient expression of MHCII_TagRFP_ containing the HLA-DRB1*1501 β-chain the sequence was amplified and cloned into the pCAGGS vector using NotI and XhoI restriction sites. The sequences encoding MHCII_TagRFP_ comprising the HLA-DRB1*0101 β-chain (HM067843.1), which is compatible with CH7C17 Jurkat T cells, was synthesized (Azenta Life Science) and cloned into pCAGGS. The correspondent mutant MHCII was generated as described above. Expression plasmid coding for a BlaM1 fusion protein^40^ were kindly provided by Silke Stertz.

### MHCII surface expression

MDCK-II cells stably expressing MHCII_mEos_ or MHCII_mEosmut_ were seeded in a 12-well plate format. The next day, cells were washed, trypsinized and centrifuged at 1300 rpm for 3 min. Cells were then resuspended and incubated with monoclonal anti-HLADR (327002, BioLegend, USA, 1:500) for 30 min on ice. Cells were washed and incubated with APC-labeled anti-mouse (BD Biosciences, 550826, 1:200) for 30 min on ice. Cells were analyzed with a BD LSRFortessa™ flow cytometer (BD Biosciences).

### Susceptibility of MDCK-II cells to infection with H18N11

MHCII_mEos_ or MHCII_mEosmut_ cells were seeded in a 24 well format and infected at an MOI of 5 for 1 h at 37°C. The inoculum was replaced with growth medium and cells were incubated for 24 hours at 37°C. After incubation, cells were washed, trypsinized and fixed with 4% PFA. Cells were then washed and stained with monoclonal anti-H18 (mouse, produced in-house; 1:100) for 30 min on ice. After washing, cells were stained with and APC-labeled anti-mouse (BD Biosciences, 550826 - 1:200) for 30 min on ice and analyzed with BD LSRFortessa™ flow cytometer (BD Biosciences)

### T cell activation

8.4×10^5^ cells BHK-21were seeded in 6-well format and transfected with 1µg pCAGGS expression vectors encoding MHCII_TagRFP_ or MHCII_TagRFPmut_ comprising the HLA-DRB1*0101 β-chain using Lipofectamine 2000 (Thermo Fisher, Germany). MHCII_TagRFP_ comprising the HLA-DRB1*1501 β-chain was transfected as negative control. The next day, 5×10^4^ transfected cells were transferred into a well of a 96-well plate and cocultured with 10^5^ CH7C17 Jurkat T cells in RPMI 1640 (Gibco, USA) supplemented with 10% FCS and 5% HEPES for 6 h at 37°C. Subsequently, cells were stained with FITC-labeled anti-CD3 antibody (BioLegend, USA, 1:200) and APC-labeled anti-CD69 antibody (Life Technologies, USA, 1:200) and analyzed with a BD FACSCanto II (BD Biosciences) flow cytometer.

### Polykaryon formation assay

8.4×10^5^ HEK293T cells were seeded in 6-well plates and cotransfected with 2 μg of pCAGGS-GFP and either pCAGGS-EV (empty vector) or pCAGGS-H18. For transfection, Lipofectamine 2000 (Thermo Fisher, Germany) was used at a DNA-to-Lipofectamine ratio of 1:2 in Opti-MEM. At 24h post-transfection, 10^5^ transfected HEK293T cells and 10^5^ MDCK-II cells stably expressing MHCII_tagRFP_ or MHCII_tagRFPmut_, were coseeded in collagen-coated 24-well plates containing growth medium (DMEM, 10% FCS, 100 U per ml penicillin, and 100 mg per ml streptomycin) and incubated at 37°C and 5% CO_2_. The following day, cells were treated with TPCK trypsin (10 μg per ml in Opti-MEM) for 30 min at 37°C. Cells were subsequently washed with PBS, exposed to pH 5 PBS for 20 min at 37°C and 5% CO_2_, and then incubated in growth medium for 2 h at 37°C and 5% CO_2_. Finally, cells were washed with PBS, fixed with 4% PFA for 20 min, and nuclei were stained in the dark for 1 h using DAPI (1:10.000) in PBS. Fluorescence images were acquired using a Zeiss Observer.ZI inverted epifluorescence microscope (Carl Zeiss) equipped with an AxioCamMR3 camera using a 20× objective.

### Production of soluble MHCII

sMHCII and sMHCII_mut_ were expressed in Expi293F cells by transfecting 100 µg of plasmid (1:1 ratio of α-chain coding plasmid: β-chain coding plasmid) for 100 ml of cells at 2×10^6^ cells per ml. Transfection was performed using the ExpiFectamine 293 Transfection Kit (Thermo Fisher Scientific) following the manufacturer instructions. 6 days later, cells were centrifuged, and the supernatant was filtered with a 0.45 µm filter (Sartorius). sMHCII molecules were purified with affinity chromatography using HisPur Cobalt Resin (Thermo Fischer Scientific) and Strep-TactinXT 4Flow high capacity resin (IBA Lifesciences), followed by size exclusion chromatography on a Superdex 200 Increase column into a buffer containing 20 mM Tris pH 8.0 and 150 mM NaCl.

### Coomassie gel staining

Proteins were separated using SDS-PAGE on a 10% polyacrylamide gel. After electrophoresis, the gel was rinsed briefly with deionized water. The gel was stained with Blauer Jonas (BIOZOL) and incubated for 1 h at room temperature. Following staining, the gel was washed with water to remove the residual solution and was imaged using Vilber E-Box.

### MHCII competition assay

MDCK-II cells stably expressing MHCII_mEos_ were seeded in a 24-well plate format. The next day, sMHCII and sMHCII_mut_ were diluted in MEM medium (Gibco, USA) supplemented with 2% FCS, mixed with 50 focus-forming units of H18N11 (rP11) and incubated at 37°C for 1 h. Cells were pre-treated with PBS containing 15 µg per ml DEAE-Dextran for 15 min. Subsequently, cells were washed, infected with the H18N11:MHCII mix and incubated for 1h at 37°C. Cells were then covered with 500 µL of overlay (DMEM, 20 mM HEPES pH 7.5, 200 µg per ml bovine serum albumin (BSA), 20 µg per ml DEAE-Dextran, 100 µg per ml NaHCO_3_, 40 µg per ml oxoid agar) and incubated at 37°C for 36 h. After overlay removal, cells were fixed with 4% PFA for 20 min, permeabilized with PBS, 0.1% Triton and blocked with PBS containing 5% FCS. Cells were then stained with a monoclonal anti-H18N11-NP antibody (produced in-house, 1:500) for 1 h at room temperature, washed and incubated with a secondary HRP-conjugated anti-mouse antibody (Dianova, 1:500) for 1 h at room temperature. After washing, focus forming units were visualized by adding the substrate solution (PBS supplemented with 0.5 µg per ml 3, 3’-diaminobenzidine (DAB), 0.5 µg per ml nickel ammonium sulfate and 0.015 % H_2_O_2_).

### Quantification of MHCII nanoclustering via PALM in fixed cells

To determine the nanoscale organization of MHCII within the plasma membrane, MDCKII cells stably expressing MHCII_mEos_ or MHCII_mEosmut_ were grown on 25 mm glass cover slips overnight, then fixed in PFA 4% for 15 min at room temperature. Glass cover slips were mounted in an AttoFluor cell chamber (ThermoFisher), for imaging. PALM was performed on a Nikon Ti Eclipse NSTORM microscope equipped with three laser lines at 405 nm (Coherent), 488 nm (Sapphire, Coherent) and a 561 nm (Sapphire, Coherent). The sample was observed through a Nikon Apo TIRF 100x Oil DIC N2 NA 1.49 objective and emitted light was detected on an Andor iXon3 DU-897 EMCCD camera (Oxford Instruments). Emitted light was directed onto the camera using a dichroic mirror and filtered through a Bright Line HC 609/64 emission filter. We typically took 20k frames at 30 ms integration time using the NIS elements software. Single emitter positions were localized using the software DECODE, the processed using custom MatLab (MathWorks) routines as described previously^21,41^.To then investigate the MHCII clustering in the presence of the virus, MHCII_mEos_ or MHCII_mEosmut_ were incubated on ice for 1 h with H18N11, at an MOI of 200. Unbound viruses were washed away with cold PBS and the cells were fixed with PFA 4% for 15 min at room temperature. For the staining, the cells were incubated for 1 h in blocking buffer (0.2% BSA in PBS). Primary polyclonal anti-H18 (rabbit, produced in-house) was diluted 1:750 in blocking buffer and the cells were incubated for 1h at room temperature. The cells were then washed 3 times in PBS and incubated for 1h at room temperature with secondary antibody anti-rabbit AlexaFluor647 (ThermoFisher, 1:250). Finally, the cells were washed 3 times in PBS. Images were acquired as described above (20k frames per acquisition).

### Virus tracking

4.5×10^5^ MHCII_TagRFP_ or MHCII_TagRFPmut_ were seeded on a 35 mm² dish (Ibidi). The next day, cells were pre-incubated for 15 min on ice. Growth medium was discarded and replaced with infection medium containing DiO (ThermoFisher, V22886)-labeled H18N11 filtered using a 0.2 µm pore size sterile filter. Cells were incubated with virus on ice for 30 minutes. Cells were then washed with cold PBS and covered with pre-warmed growth medium. The samples were imaged at 37° and 5% CO_2_ using a Nikon CSU-W1 spinning disk confocal microscope for 20 min.

### Measurement of MHCII single protein diffusion using sptPALM in live cells

H18N11 virus was diluted 1:1 in PBS and labeled with DiD (ThermoFisher, V22887) for 30 min at room temperature in the dark. Viral particles were separated from unbound dye by gel filtration (Nap5 columns, equilibrated with PBS) (Cytiva, 17085201). The labeled virus particles were then vortexed, spun down and filtered over a 0.22 µm pore size filter and subsequently resuspended in PBS. 3D-Epoxy Glass Coverslips (PolyAN, 10400206) were mounted at the bottom of a 6-well chamber (ibidi, 80828). Primary buffer (150 mM Na_2_HPO_4_, 50 mM NaCl, pH 8.5) was distributed in the wells and the labeled virus was added and incubated overnight at 4°C. The following morning, the virus was discarded and the secondary buffer (50 mM ethanolamine, 100 mM Tris, pH 9.0) was added for 1h at room temperature. Wells were then washed first with PBS and then with infection medium (DMEM, 0.2% BSA, penicillin-streptomycin). 3×10^4^ MHCII_mEos_ or MHCII_mEosmut_ cells were seeded in each well and cultured until fully attached, typically overnight. MHCII molecules were analyzed via PALM as described above but at 37°C and 5% CO_2_. Under live cell conditions, we typically acquired 10k frames at 30 ms integration time using the NIS elements software.

Single emitter positions were localized using the software DECODE^41^ and trajectories were reconstructed using the TrackPy Python library. Localizations were linked using a KDTree algorithm with a maximum search range of 0.8 pixels and a zero-frame-gap memory. The adaptive stop for solving oversized subnets was set to 0.1 with a step size multiplicator of 0.9. For further analysis, only tracks with a minimum length of 10 frames were considered. Ensemble drift xy(t) was calculated and subtracted from the tracks. Mean squared displacement (MSD, equation 1) was determined and plotted individually for all molecules. For all tracks, the initial diffusion coefficients (D_1-3_) and the power law exponent alpha were calculated by applying a linear fit in log space (Equation 2). Power law exponents and diffusion coefficients for virus spots were calculated within an 800×800 nm square around an immobilized virus.

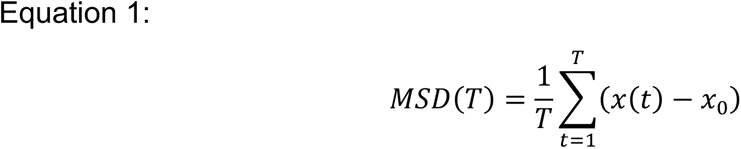

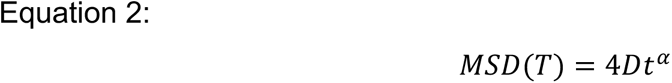

---

**Supplementary Figure 1: Validation of fluorescently-labeled MHCII. (A)** HEK293T cells were transiently transfected with MHCII_mEos_ and exposed for 20 min under UV light (365 nm) to test for photoconversion of the mEos3.2 fluorophore. Scale bar 100 µm. **(B)** Schematic representation of the fluorescently labeled human MHCII constructs (HLA-DR): MHCII_TagRFP_ and MHCII_TagRFPmut_. The Influenza HA307-319 peptide was fused to the N-terminus of the beta chain. The red fluorescent TagRFP was fused to the C-terminus of the MHCII beta chain. MHCII_TagRFP_ comprises an 11 amino acid substitution in the alpha2 subunit (blue). **(C)** Infection rate of MDCKII stably expressing MHCII_TagRFP_ or MHCII_TagRFPmut_ at 24 hours post-infection with H18N11 determined by flow cytometry. **(D)** Infection rate of HEK293T cells transiently transfected with MHCII_TagRFP_ and MHCII_TagRFPmut_ of the indicated haplotype determined at 24 hours post infection with H18N11 determined by flow cytometry. **(E)** pH-induced polykaryon formation of HEK293T cells transiently expressing empty vector and GFP with MDCK-II stably expressing MHCII_TagRFP_ or MHCII_TagRFPmut_. Images are representatives of 3 independent experiments. Scale bar 100 µm.

**Supplementary Figure 2: MHCII_mEos_ and MHCII_mEosmut_ organize and behave similarly when expressed in MDCK-II cells.** To explore if MHCII_mEos_ and MHCII_mEosmut_ are differently organized in uninfected MDCK-II cells, we imaged the apical surface of fixed cells and the basolateral surface of live cells using PALM. Imaging fixed cells (**A**) reveals a similar cluster organization of MHCII_mEosmut_ as shown for MHCII_mEos_ in Fig. 3B. The cluster size quantification is shown in Fig. 3C. sptPALM of MHCII_mEos_ and MHCII_mEosmut_ revealed similar diffusion coefficients (**B**) and organization of mobile and immobile proteins (**C-H**). (C and F) show a rendering of a live cell sptPALM acquisition of MHCII_mEos_ (wt) and MHCII_mEosmut_ (mut), respectively, revealing clusters as observed in fixed cells (A, and Fig. 3B). D and G show the highlighted areas in (C and F). **(E and H)** To visualize the distribution of mobile and immobile receptors, trajectories with D<0.01 µm^2^/s were considered as immobile and colored in red. Trajectories of mobile receptors are shown in green. Again, a similar distribution can be observed of MHCII_mEos_ and MHCII_mEosmut_. Scale bars: A, 1 µm; C, 1.3 µm; F, 1.2 µm.

## Notes

### Competing Interest Statement

The authors have declared no competing interest.

## References

1 Fouchier Ron, A. M., et al. Characterization of a Novel Influenza A Virus Hemagglutinin Subtype (H16) Obtained from Black-Headed Gulls. Journal of Virology 79, 2814–2822, doi:10.1128/jvi.79.5.2814-2822.2005 (2005).

2 Kida, H. & Yanagawa, R. Isolation and characterization of influenza a viruses from wild free-flying ducks in Hokkaido, Japan. Zentralbl Bakteriol Orig A 244, 135–143 (1979).

3 Webster, R. G., Bean, W. J., Gorman, O. T., Chambers, T. M. & Kawaoka, Y. Evolution and ecology of influenza A viruses. Microbiological Reviews 56, 152–179, doi:10.1128/mr.56.1.152-179.1992 (1992).

4 Tong, S. et al. A distinct lineage of influenza A virus from bats. Proceedings of the National Academy of Sciences 109, 4269–4274, doi:10.1073/pnas.1116200109 (2012).

5 Tong, S. et al. New World Bats Harbor Diverse Influenza A Viruses. PLOS Pathogens 9, e1003657, doi:10.1371/journal.ppat.1003657 (2013).

6 Campos, A. C. A. et al. Bat Influenza A(HL18NL11) Virus in Fruit Bats, Brazil. Emerging Infectious Disease journal 25, 333, doi:10.3201/eid2502.181246 (2019).

7 Li, Q. et al. Structural and functional characterization of neuraminidase-like molecule N10 derived from bat influenza A virus. Proceedings of the National Academy of Sciences 109, 18897–18902, doi:10.1073/pnas.1211037109 (2012).

8 Sun, X. et al. Bat-Derived Influenza Hemagglutinin H17 Does Not Bind Canonical Avian or Human Receptors and Most Likely Uses a Unique Entry Mechanism. Cell Reports 3, 769–778, 10.1016/j.celrep.2013.01.025 (2013).

9 Zhu, X. et al. Hemagglutinin homologue from H17N10 bat influenza virus exhibits divergent receptor-binding and pH-dependent fusion activities. Proceedings of the National Academy of Sciences 110, 1458–1463, doi:10.1073/pnas.1218509110 (2013).

10 Moreira, É. A. et al. Synthetically derived bat influenza A-like viruses reveal a cell type-but not species-specific tropism. Proceedings of the National Academy of Sciences 113, 12797–12802, doi:10.1073/pnas.1608821113 (2016).

11 Ciminski, K. et al. Bat influenza viruses transmit among bats but are poorly adapted to non-bat species. Nature Microbiology 4, 2298–2309, doi:10.1038/s41564-019-0556-9 (2019).

12 Karakus, U. et al. MHC class II proteins mediate cross-species entry of bat influenza viruses. Nature 567, 109–112, doi:10.1038/s41586-019-0955-3 (2019).

13 Giotis, E. S. et al. Entry of the bat influenza H17N10 virus into mammalian cells is enabled by the MHC class II HLA-DR receptor. Nat Microbiol 4, 2035–2038, doi:10.1038/s41564-019-0517-3 (2019).

14 Brown, J. H. et al. Three-dimensional structure of the human class II histocompatibility antigen HLA-DR1. Nature 364, 33–39, doi:10.1038/364033a0 (1993).

15 Rock, K. L., Reits, E. & Neenes, J. Present Yourself! By MHC Class I and MHC Class II Molecules. Trends in Immunology 37, 724–737, 10.1016/j.it.2016.08.010 (2016).

16 Olajide, O. M. et al. Evolutionarily conserved amino acids in MHC-II mediate bat influenza A virus entry into human cells. PLOS Biology 21, e3002182, doi:10.1371/journal.pbio.3002182 (2023).

17 Sauter, N. K. et al. Hemagglutinins from two influenza virus variants bind to sialic acid derivatives with millimolar dissociation constants: a 500-MHz proton nuclear magnetic resonance study. Biochemistry 28, 8388–8396, doi:10.1021/bi00447a018 (1989).

18 Xiong, X. et al. Receptor binding by an H7N9 influenza virus from humans. Nature 499, 496–499, doi:10.1038/nature12372 (2013).

19 Vachieri, S. G. et al. Receptor binding by H10 influenza viruses. Nature 511, 475–477, doi:10.1038/nature13443 (2014).

20 Weis, W. et al. Structure of the influenza virus haemagglutinin complexed with its receptor, sialic acid. Nature 333, 426–431, doi:10.1038/333426a0 (1988).

21 Sieben, C., Sezgin, E., Eggeling, C. & Manley, S. Influenza A viruses use multivalent sialic acid clusters for cell binding and receptor activation. PLOS Pathogens 16, e1008656, doi:10.1371/journal.ppat.1008656 (2020).

22 Overeem, N. J., van der Vries, E. & Huskens, J. A Dynamic, Supramolecular View on the Multivalent Interaction between Influenza Virus and Host Cell. Small 17, 2007214, 10.1002/smll.202007214 (2021).

23 Anderson, H. A., Hiltbold, E. M. & Roche, P. A. Concentration of MHC class II molecules in lipid rafts facilitates antigen presentation. Nature Immunology 1, 156–162, doi:10.1038/77842 (2000).

24 de la Fuente, H. et al. Synaptic clusters of MHC class II molecules induced on DCs by adhesion molecule-mediated initial T-cell scanning. Mol Biol Cell 16, 3314–3322, doi:10.1091/mbc.e05-01-0005 (2005).

25 Kropshofer, H. et al. Tetraspan microdomains distinct from lipid rafts enrich select peptide–MHC class II complexes. Nature Immunology 3, 61–68, doi:10.1038/ni750 (2002).

26 Betzig, E. et al. Imaging Intracellular Fluorescent Proteins at Nanometer Resolution. Science 313, 1642–1645, doi:doi:10.1126/science.1127344 (2006).

27 Hess, S. T., Girirajan, T. P. & Mason, M. D. Ultra-high resolution imaging by fluorescence photoactivation localization microscopy. Biophys J 91, 4258–4272, doi:10.1529/biophysj.106.091116 (2006).

28 Zhang, M. et al. Rational design of true monomeric and bright photoactivatable fluorescent proteins. Nature Methods 9, 727–729, doi:10.1038/nmeth.2021 (2012).

29 Cuellar-Camacho, J. L. et al. Quantification of Multivalent Interactions between Sialic Acid and Influenza A Virus Spike Proteins by Single-Molecule Force Spectroscopy. Journal of the American Chemical Society 142, 12181–12192, doi:10.1021/jacs.0c02852 (2020).

30 Bosch, B., Heipertz, E. L., Drake, J. R. & Roche, P. A. Major histocompatibility complex (MHC) class II-peptide complexes arrive at the plasma membrane in cholesterol-rich microclusters. J Biol Chem 288, 13236–13242, doi:10.1074/jbc.M112.442640 (2013).

31 Qian, H., Sheetz, M. P. & Elson, E. L. Single particle tracking. Analysis of diffusion and flow in two-dimensional systems. Biophysical Journal 60, 910–921, 10.1016/S0006-3495(91)82125-7 (1991).

32 Masaki, K. et al. Ligation of MHC Class II Induces PKC-Dependent Clathrin-Mediated Endocytosis of MHC Class II. Cells 9 (2020).

33 Furuta, K., Ishido, S. & Roche, P. A. Encounter with antigen-specific primed CD4 T cells promotes MHC class II degradation in dendritic cells. Proceedings of the National Academy of Sciences 109, 19380–19385, doi:10.1073/pnas.1213868109 (2012).

34 Andreae, S., Buisson, S. & Triebel, F. d. r. MHC class II signal transduction in human dendritic cells induced by a natural ligand, the LAG-3 protein (CD223). Blood 102, 2130–2137, doi:10.1182/blood-2003-01-0273 (2003).

35 Huby, R. D. J., Dearman, R. J. & Kimber, I. Intracellular Phosphotyrosine Induction by Major Histocompatibility Complex Class II Requires Co-aggregation with Membrane Rafts*. Journal of Biological Chemistry 274, 22591–22596, 10.1074/jbc.274.32.22591 (1999).

36 Eierhoff, T., Hrincius, E. R., Rescher, U., Ludwig, S. & Ehrhardt, C. The Epidermal Growth Factor Receptor (EGFR) Promotes Uptake of Influenza A Viruses (IAV) into Host Cells. PLOS Pathogens 6, e1001099, doi:10.1371/journal.ppat.1001099 (2010).

37 White, J., Kartenbeck, J. & Helenius, A. Membrane fusion activity of influenza virus. The EMBO Journal 1, 217–222-222, 10.1002/j.1460-2075.1982.tb01150.x (1982).

38 Ciminski, K., Pfaff, F., Beer, M. & Schwemmle, M. Bats reveal the true power of influenza A virus adaptability. PLOS Pathogens 16, e1008384, doi:10.1371/journal.ppat.1008384 (2020).

39 Kessler, S. et al. Deciphering bat influenza H18N11 infection dynamics in male Jamaican fruit bats on a single-cell level. Nature Communications 15, 4500, doi:10.1038/s41467-024-48934-6 (2024).

40 Karakus, U. et al. H19 influenza A virus exhibits species-specific MHC class II receptor usage. Cell Host & Microbe, doi:10.1016/j.chom.2024.05.018 (2024).

41 Speiser, A. et al. Deep learning enables fast and dense single-molecule localization with high accuracy. Nature Methods 18, 1082–1090, doi:10.1038/s41592-021-01236-x (2021).

