## Supplementary figures and images for "The bat Influenza A virus subtype H18N11 induces nanoscale MHCII clustering upon host cell attachment"

### Figure SI 1

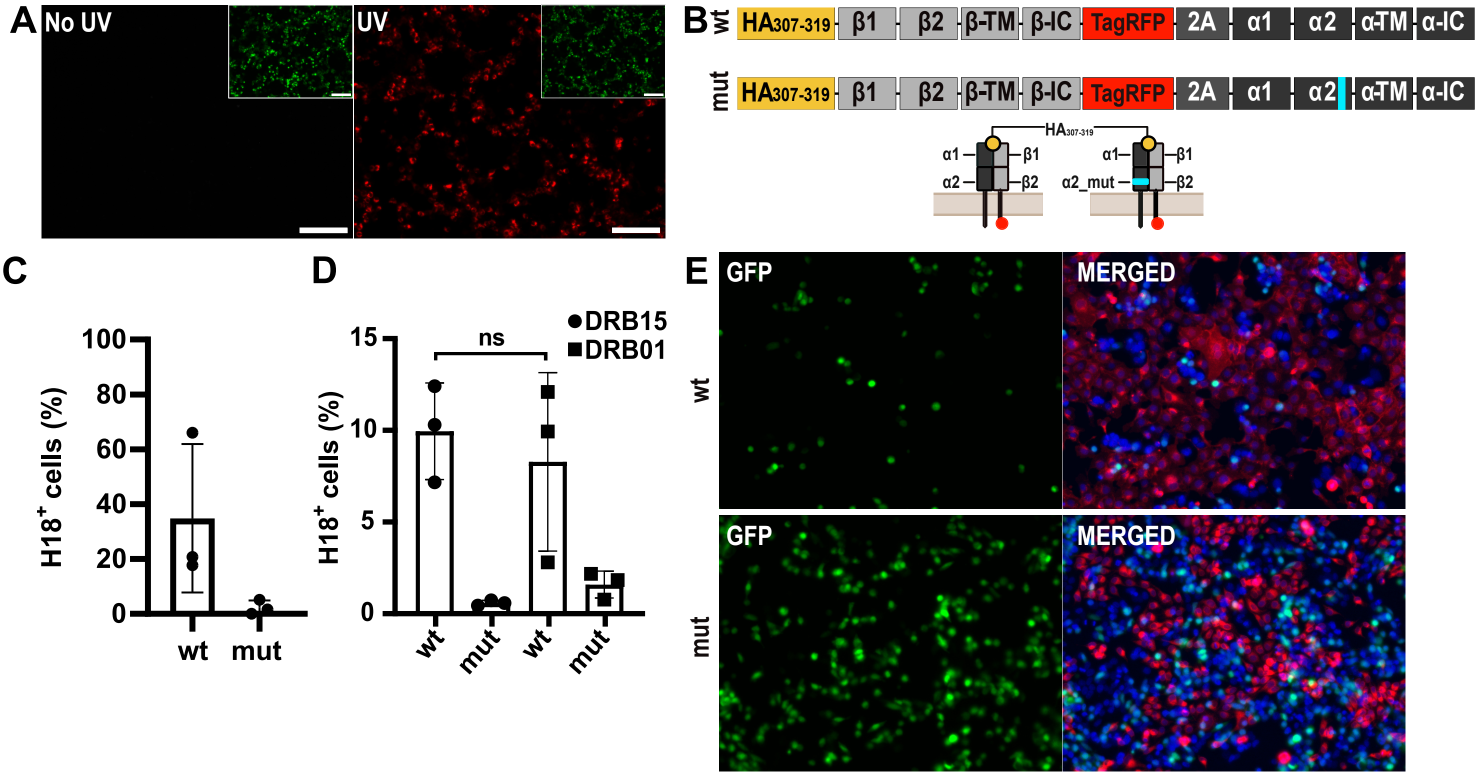

### Figure SI 2

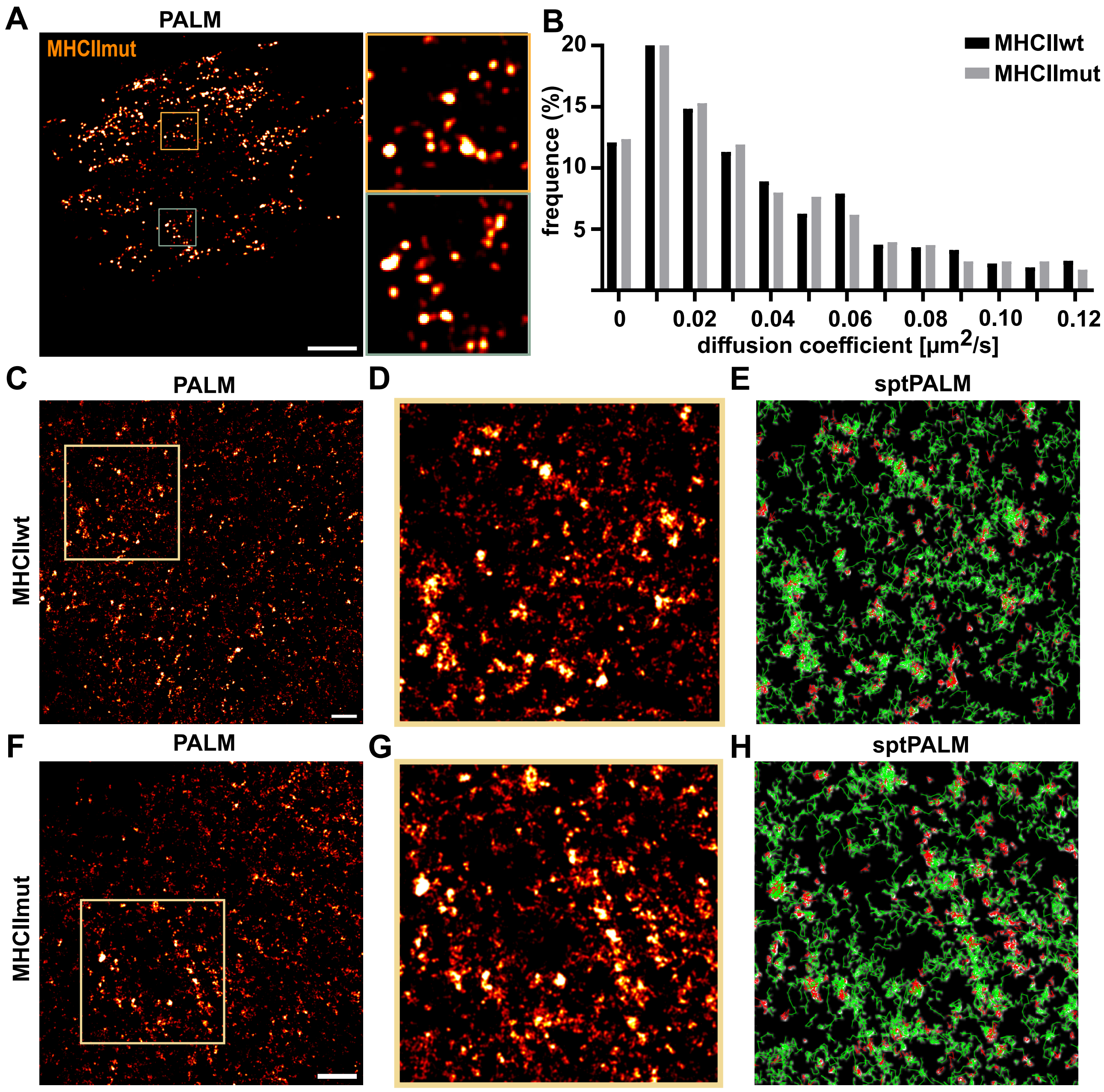
